# Sequence conservation and structural features that are common within TRP channels

**DOI:** 10.1101/2021.09.06.459056

**Authors:** Deny Cabezas-Bratesco, Charlotte K. Colenso, Katina Zavala, Danielle Granata, Vincenzo Carnevale, Juan C. Opazo, Sebastian E. Brauchi

**Affiliations:** Instituto de Fisiologia, Facultad de Medicina, Universidad Austral de Chile, Valdivia, 511-0566, Chile; School of Cellular and Molecular Medicine, University of Bristol, Bristol, BS8 1TD, UK; Instituto de Ciencias Ambientales y Evolutivas, Facultad de Ciencias, Universidad Austral de Chile, Valdivia, Chile; Institute for Computational Molecular Science, Temple University, Philadelphia, USA; Integrative Biology Group, Universidad Austral de Chile, Valdivia, Chile; Millennium Nucleus of Ion Channel-associated Diseases (MiNICAD), Valdivia, Chile; Janelia Research Campus, Howard Hughes Medical Institute, Ashburn, VA, 20147, USA

**Author notes:** D.C-B. and C.K.C. equally contributed to this work. Correspondence should be addressed to Dr. Sebastian Brauchi.

**Keywords:** MSA, evolution, allosterism, structure

## Abstract

TRP proteins are a large family of cation selective channels, surpassed in variety only by voltage-gated potassium channels. Detailed molecular mechanisms governing how membrane voltage, ligand binding, or temperature can induce conformational changes promoting the open state of the channel are still missing for TRP channels. Aiming to unveil distinctive structural features common to the transmembrane domains within the TRP family, we performed bioinformatic analyses over a large set of TRP channel genes. Here we report a discrete and exceptionally conserved set of residues. This fingerprint is composed of eleven residues localized at equivalent three-dimensional positions in TRP channels from the different subtypes. Moreover, these amino acids are arranged in three groups, connected by a set of aromatics located at the core of the transmembrane structure. We hypothesize that differences in the connectivity between these different groups of residues harbors the apparent differences in coupling strategies used by TRP subgroups.

## INTRODUCTION

Transient receptor potential (TRP) proteins constitute a large family of cation-selective ion channels involved in a number of physiological functions (Clapham, 2003; Nilius *et al.,* 2007). TRPs have been related to all voltage-gated cation channels (VGCC), two-pore sodium channels (TPCNs), and CatSper channels (Yu *et al.,* 2005). The TRP family in vertebrates is composed by of 28 channels, belonging to two major groups and seven subfamilies or subtypes: TRPA1, TRPV, TRPC, TRPM, TRPN, PKD2s, and TRPML (Ramsey *et al.,* 2006). While group I gathers members of TRPC, TRPM, TRPV, TRPA, and TRPN subfamilies, group II is composed of members of the TRPML and PKD2 channels (Venkatachalam 2007, Himmel 2020). In general, TRP channels share poor cation selectivity and a loose sequence similarity (Ramsey *et al.,* 2006). Cataloged as polymodal ion channels, they have the ability to integrate multiple stimuli (e.g., chemical, mechanical, electrical, and thermal) to promote channel opening. Such polymodality observed in TRPs has been explained in terms of allosteric interactions (Brauchi *et al.,* 2004; Latorre *et al.,* 2009). Different lines of research have shown that sensor modules-activated by specific mechanisms-couple to each other and to the channel pore, modulating permeation (Hui *et al.,* 2003; Castillo *et al.,* 2018; Zubcevic *et al.,* 2019; Zhao *et al.,* 2020; Yang *et al.,* 2018; Yang *et al.,* 2020).

Structural data revealed that TRP channels share the general architecture of VGCCs (Kasimova *et al.,* 2016; Cheng, 2018; Cao, 2020). They assemble as domain-swapped homotetramers, with monomers containing six transmembrane segments (*i.e.*, TM1-TM6) flanked by cytoplasmic N- and C-terminal domains. The transmembrane helices TM5 and TM6, together with the section between them, give shape to the conductive pore (Ramsey *et al.,* 2006; Liao *et al.,* 2013). The first four transmembrane helices and part of the intracellular domains have been described as regulatory regions as they provide binding sites for agonists and cofactors (Steinberg *et al.,* 2014; Voolstra & Huber, 2014; Cao, 2020). The transmembrane region between TM1 and TM4 shows fundamental differences with VGCCs, where the absence of charged residues within the transmembrane region should be underscored (Palovcak *et al.,* 2015; Cao, 2020). This could explain the modest voltage dependence observed in TRP channels (TRPMs and TRPVs are tenfold less voltage dependent compared to VGCCs) that might be supported by residues located in the pore region (Liu *et al.,* 2009; Yang *et al.,* 2020). The transmembrane region between TM1 and TM4 hosts binding pockets for ligands in all the different TRP channel subtypes, serving as a ligand-binding domain (LBD) in TRPs (Steinberg *et al.,* 2014; Huffer *et al.,* 2020). Nevertheless, this LBD has been historically referred to as the voltage-sensing like domain (VSLD).

Regardless of the considerable variability in sequence similarity and physiological function, it is clear that TRP channels are closely related, specially those that belongs to group I (Kadowaki, 2015; Peng *et al.,* 2015; Arias-Darraz *et al.,* 2015). The structural understanding of TRP proteins has been strengthened by technical advances allowing the recent release of a large set of high-resolution cryo-electron microscopy (cryo-EM) structures (Cao, 2020; Samanta *et al.,* 2019). Led by the structure of TRPV1, nowadays structures are available for at least one member of each subfamily, including crTRP1 from the green algae *Chlamydomonas reinhardtii* (McGoldrick *et al.,* 2019; Cao, 2020). Together with advances in structural biology, intense research during recent years has been focused into understanding the molecular mechanisms supporting TRP activation and regulation (Hofmann *et al.,* 2017; Yang & Zheng, 2017; Yang *et al.,* 2018; Singh, *et al.,* 2018; Hilton *et al.,* 2019; Zhang *et al.,* 2019; Zubcevic *et al.,* 2019; Zhao *et al.,* 2020, Nadezhdin *et al.,* 2021; Nadezhdin *et al.,* 2021a). Although a consistent picture accounting for general principles governing TRP channels’ mechanics is still missing, several structural features have been identified of great importance. Among these the N- and C-terminal domains flanking the transmembrane region namely the preTM1 and the TRP domain helix (TDh) respectively. These elements, seemingly related to the integration of molecular mechanics during activation, are present in all members of the group I of TRPs and absent in TRP channels from group II (McGoldrick *et al.,* 2019; Zhao *et al.,* 2020; Nadezhdin *et al.,* 2021).

Here we studied TRP channel proteins from a large variety of organisms aiming to find conserved amino acid motifs. We found a discrete number of highly conserved residues in all TRP channels from group I (GI-TRPs). Moreover, strong patterns of conservation were found unique to the different TRP subtypes. The amino acid conservation can be traced down to TRP channels from unicellular organisms, suggesting a robust architectural design. In addition, we identified a group of aromatic residues facing the core of the LBD (*i.e.* TM 1-4) in all subtypes. In agreement with our phylogenetic reconstruction this aromatic core can be traced down to crTRP1, it is absent in VGCCs, and present in a rudimentary form in TPCNs. Further analyses of the highly conserved residues unveiled inter-subunit interactions connecting the aromatic core of one subunit with signature residues located at the selectivity filter of the neighboring subunit. Overall, our results suggest that TRP channel specialization has been built around the connectivity of heavily conserved distant residues that are located at critical sites within the structure.

## RESULTS

### Phylogenetic relationships of GI-TRPs

We first investigated the phylogenetic position of the unicellular TRP channels included in this study. To this end we reconstructed a phylogenetic tree including TRPs from amniotes in addition to unicellular sequences we previously identified as TRP channels (Fig. 1) (Arias-Darraz *et al.,* 2015). The topology of the phylogenetic reconstruction is in agreement with previous reports (Himmel *et al.,* 2020). According to our phylogenetic tree, we recovered TRP channels from unicellular organisms into five clades (UC1-5, Fig. 1). A clade containing TRP sequences from *Dictyostelium discoideum*, *Dictyostelium purpureum* and *Paramecium tetraurelia* was recovered sister to the clade including TRPVs and TRPA1 sequences (UC3, Fig. 1). In turn, a group including sequences from *Dictyostelium discoideum*, *Dictyostelium purpureum* and *Coccomyxa subellipsoidea* was recovered sister to the UC3/ TRPVs/TRPA1 clade (UC2, Fig. 1). A well supported clade containing five sequences from *Chlamydomonas reinhardtii*, one sequence from *Leishmania mexicana*, two sequences from *Micromonas pusilla* and a single sequence from *Volvox carteri* was recovered sister to the above mentioned clade (UC1, Fig. 1). Thus, this branching pattern suggests that these three unicellular clades are more related to TRPVs and TRPA1 channels than to any other channel in the TRP gene family. We also recovered a single sequence from *Chlamydomonas reinhardtii* (*i.e.* crTRP1) sister to the TRPC clade with strong support (UC5, Fig. 1), suggesting that this sequence from this single-cell green algae shares a common ancestor more recently in time with TRPCs than with any other gene family member.

**Figure 1.**
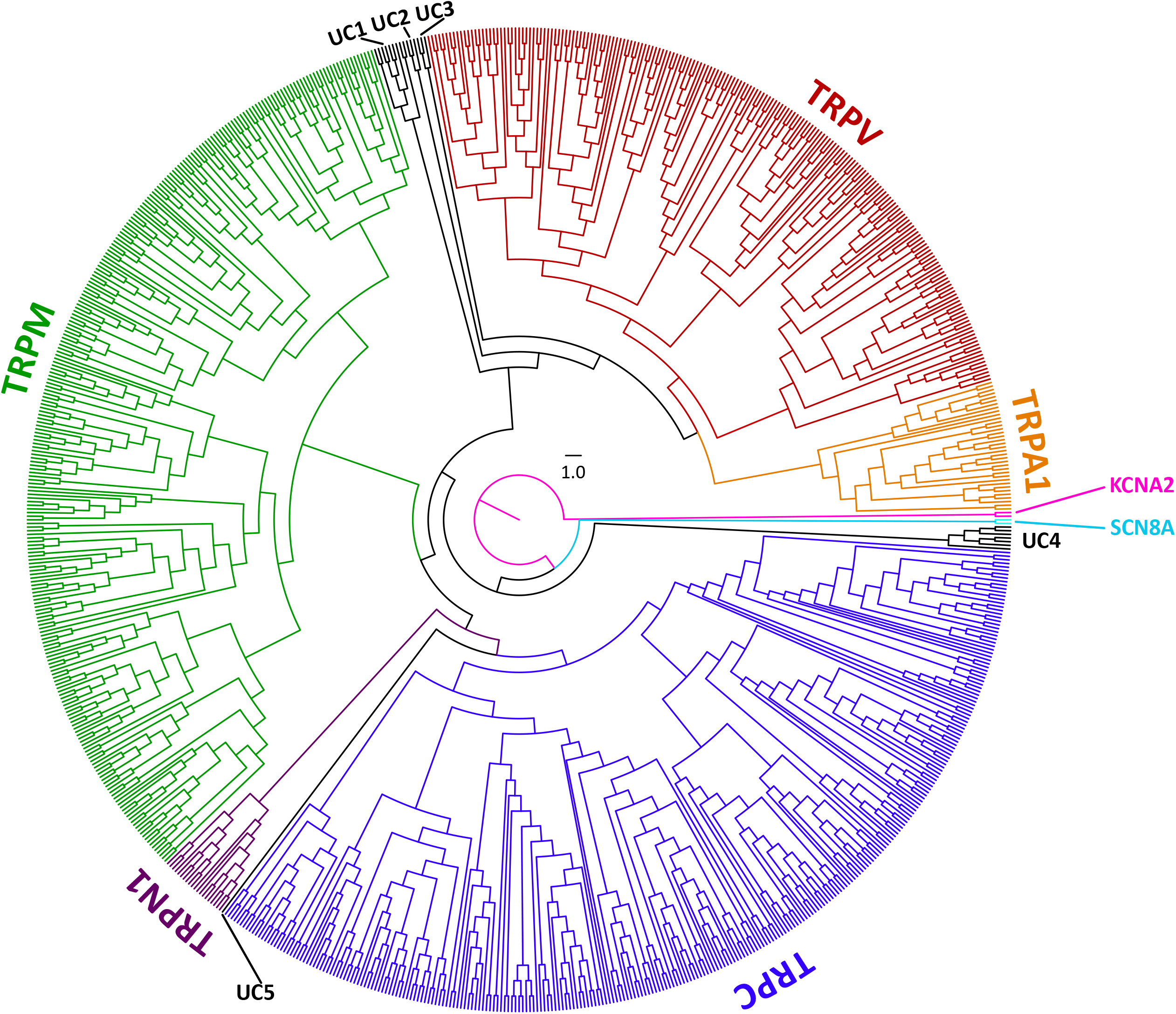
Maximum likelihood tree showing relationships among Group I of TRP channels from amniotes and the putative TRPs from unicellular organisms. The scale denotes substitutions per site and colors represent lineages. Human (*Homo sapiens*) and chicken (*Gallus gallus*) Potassium voltage-gated channel subfamily A member 2 (KCNA2), and Sodium Voltage-Gated Channel Alpha Subunit 8 (SCN8A) sequences were included as an outgroup.

Finally, we recovered a clade containing sequences from *Leishmania infantum*, *Leishmania major*, *Leishmania mexicana, Trypnosoma cruzi* and *Paramecium tetraurelia* sister to all other TRP sequences included in our study (UC4, Fig. 1).

### Multiple sequence alignments identify a discrete set of highly conserved residues

We then focused our attention on the transmembrane regions and the immediate flanking segments that are common to all TRPs. In particular, the segment containing the last portion of the pre-TM1 region, the transmembrane region, and the TRP domain helix (TDh). We then constructed a multiple sequence alignment using TRPs included in our phylogenetic analysis (including those from UC groups 1-5) and a set of non-redundant TRP channel sequences gathered from the UniProt database (Figure 2a). In agreement with previous observations (Huffer *et al.,* 2020), gaps are in general confined to extracellular loops and almost nonexistent at the intracellular linker between TM4-TM5 and the TDh (Figure 2a). Moreover, the pattern of these gaps in the extracellular loops seems to be a good predictor for the subfamily grouping. This is the case of the pore loop region, which is comparatively longer in TRPM channels, or the extended TM3-TM4 loop seen in channels from the TRPC family (Figure 2a). This observation holds true for the individual alignment of each subfamily, where a correlation between the pattern of the gaps and the different members within subfamilies is evident (Supplementary Figure 1). We tested the usefulness of these patterns as predictors by looking at *Drosophila* channels TRPL and TRPgamma that were included in the dataset pulled from Uniprot. We observed that TRPL and TRPgamma segregate to TRPCs and within the subtype, they showed a unique pattern of sequence similarity and gaps-similar to a bar code-allowing their identification from other subfamily members (Supplementary Figure 1).

**Figure 2.**
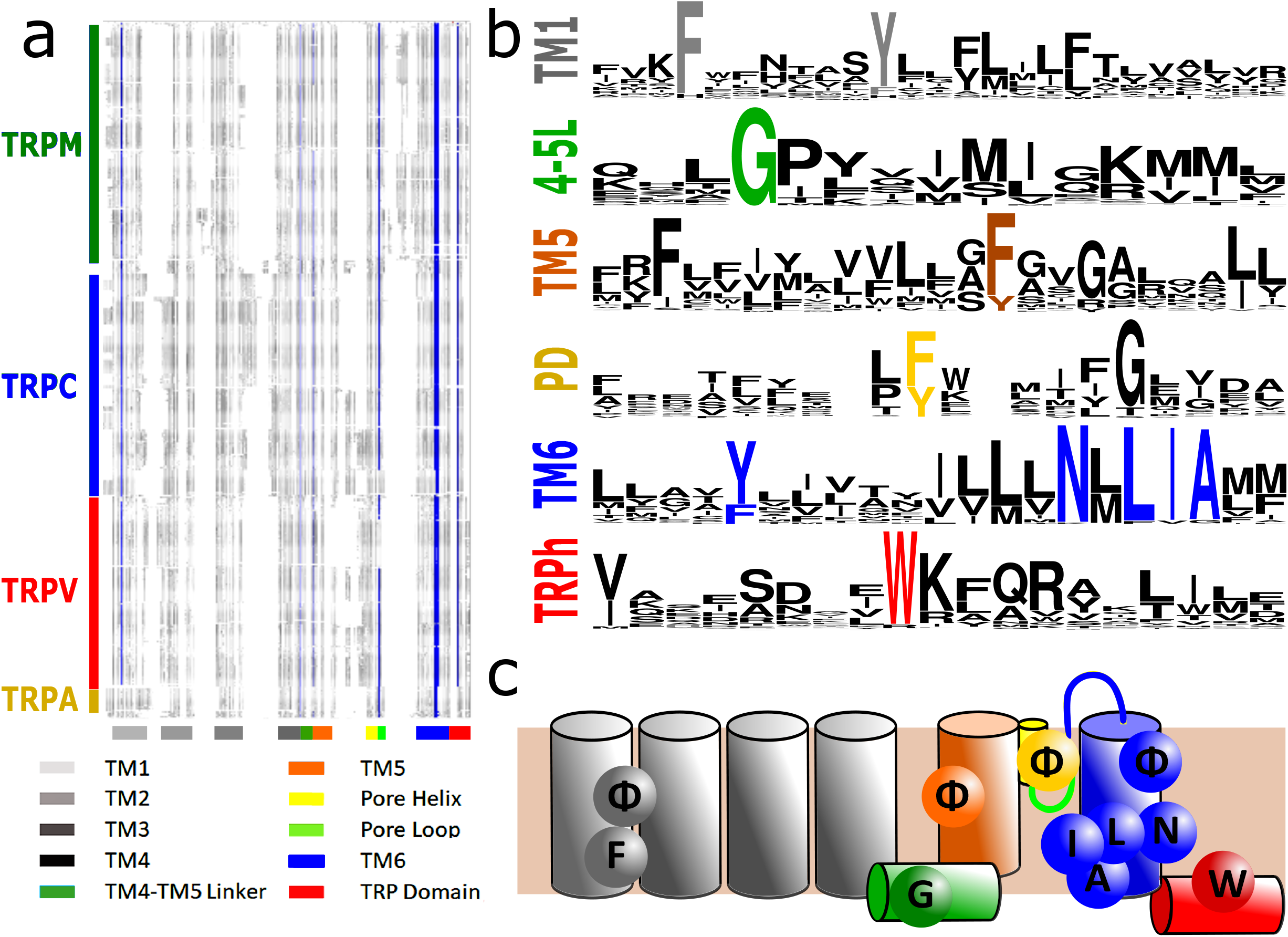
Conserved residues in CI-TRPs. (a) Schematic representation of the MSA. White blanks correspond to gaps in the alignment. Gray shades represent non-conserved residues. Blue shades correspond to sequence identities (>90%). TRP channel subfamilies are indicated on the l eft. Structural mapping is indicated below in color code. (b) Sequence logos for the TRP family, depicting highly conserved residues (>90% identity). (c) Cartoon of a TRP channel monomer depicting the location of conserved residues in the secondary structure. Φ denotes 6 carbon aromatic residues (i.e., Tyr or Phe).

A thorough analysis of the distribution of amino acids identified a discrete set of highly conserved residues (identity >90%) common to the TRP superfamily (Figure 2; Table 1). 70% of TRP channels analyzed have 11 conserved and non-contiguous amino acids we interpreted as an amino acid signature or fingerprint (**F Φ G Φ Φ Φ N L I A W**, where Φ could be either Phe or Tyr; Figure 2b). We observed that 95% of the TRP channels contained (1311 sequences in total) at least 8 of these conserved side chains. In contrast, the bacterial potassium channel KvAP shows five of these residues while Kv1.2 exhibits only three (Table 1).

**Table 1.**
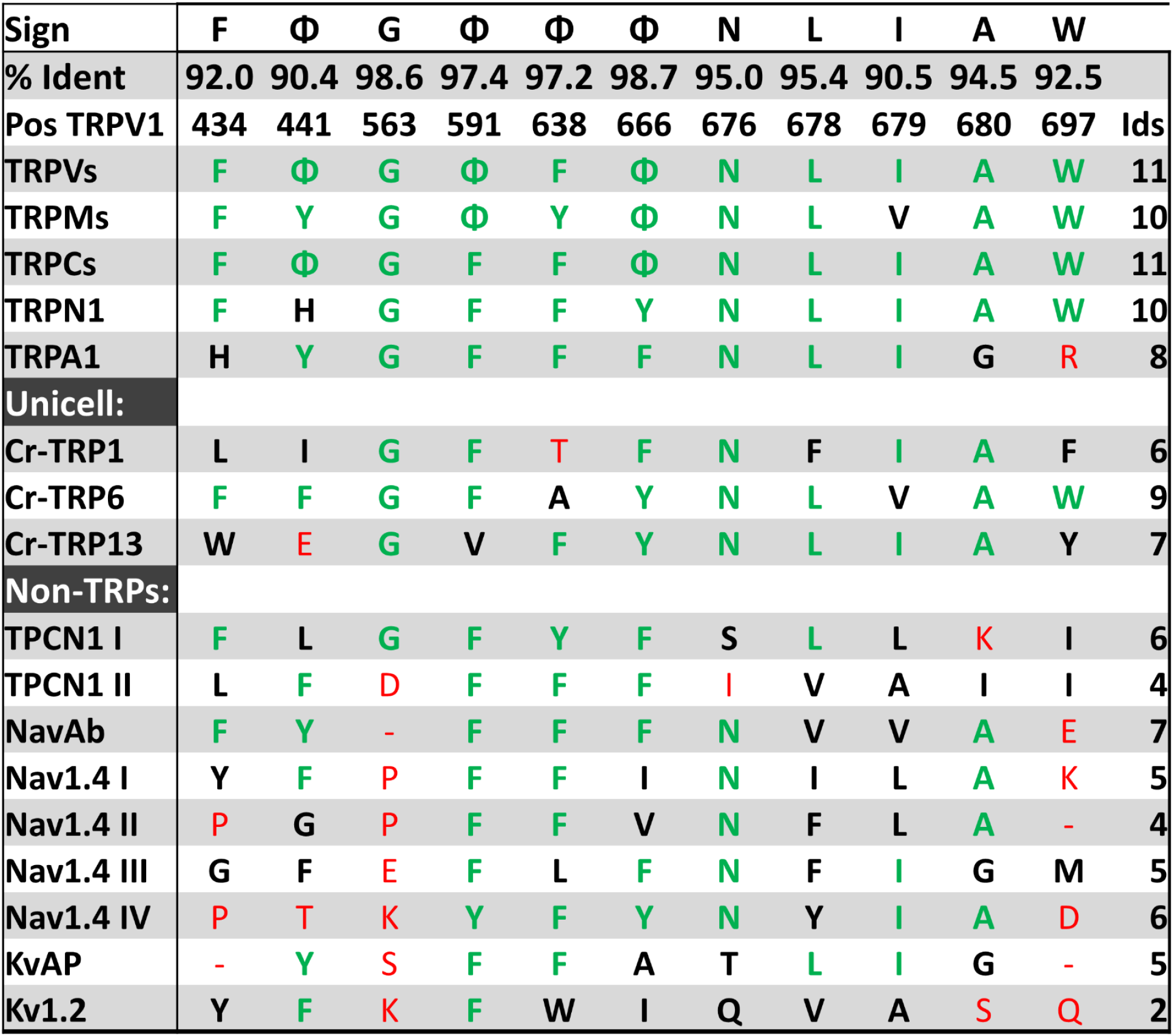
Conservation and identity of TRP fingerprint residues in different channels. Residue numbering corresponds to the rat TRPV1 sequence (poe TRPV1). TRP and non-TRP channels are presented. The number in the last column corresponds to the total number of fingerprint residues for each channel. For non-TRPs, sequence and structural alignments were used to determine equivalent positions. Green corresponds to identities while black represents homology. Red shades denote non-conserved residues. Minus signs represent gaps (non aligning residues). Φ denotes 6 carbon aromatic residues (*i.e.,* Tyr or Phe).

This fingerprint was used to compare the conservation between TRP channel subtypes. While TRPCs and TRPVs conserve the complete set of 11 residues (**F Φ G Φ F Φ N L I A W**), TRPMs and TRPNs show 10 (**F Y G Φ Y Φ N L** V **A W**), and TRPA1 only 8 (H **Y G F F F N L I** G R). The differences observed in TRPA1 can be mapped to the first turn of the TM1 helix and TDh. The latter is considered an important modulator of TRP gating and although different in sequence in TRPA1 channels when compared to other GI-TRPs, it is structurally equivalent (Paulsen *et al.,* 2015). Thus, our results suggest that such specific specialization in TRPA originated after the divergence from the common ancestor shared with TRPVs and is unlikely to be found outside this group. Within the unicellular group, crTRP1 shows only 6 identities, that increases up to 7 and 10 for the case of the unicellulars that are closer to TRPA and TRPV clades respectively (*i.e.*, UC1 to 3 groups in Figure 1; Table 1).

There are some highly conserved residues that were not considered in our analysis because they score just below the threshold. These include a highly conserved glycine (88.1%) commonly found at the selectivity filter, a phenylalanine (85.1%) at the beginning segment of TM5 and an aspartic acid (83,2%) localized at the end of TM4-TM5 linker. The latter two residues are in close proximity to L678 (according to rTRPV1 numbering) a sidechain associated to channel response to both agonist and pH in different TRPs channels (Boukalova *et al.,* 2010; Du *et al.,* 2009; Kasimova *et al.,* 2017; Klausen *et al.,* 2014).

Several groups have suggested an evolutionary relationship between TPCNs and TRP channels (Clapham & Garbers, 2005; Galione, 2010). TPCN channels are thought to be asymmetric (Penny *et al.,* 2016; Kintzer & Stroud, 2017). In support of the argument of asymmetry, while one domain (D1) shows 8 coincidences with the TRP fingerprint, the other domain (D2) is more similar to eukaryotic voltage-gated sodium channels (Nav) with only 5 coincidences. Interestingly, the monomeric bacterial channel NabAB (Payandeh *et al.,* 2011) shares seven of these eleven signature residues, and the mammalian Nav1.4 exhibits only 6 residues in domain IV, which holds the highest number of hits compared to domains I-III (Table 1). Thus, in our analysis, TPCN and Nav have the larger score of similarities in residue conservation outside the TRP family. Overall, phylogenetic and primary sequence analyses provide strong support for a fingerprint in TRP channels that is composed of 8 to 11 non-contiguous residues (**F Φ G Φ Φ Φ N L I A W**).

### Sequence conservation highlights structural features

To visualize the position of these fingerprint residues, TRP channel structures from the different families were compared (*i.e.*, TRPV1, PDBID 5IRZ; TRPM8, PDBID 6BPQ; TRPC3, PDBID 5ZBG; TRPA1, PDBID 3J9P and TRP1, PDBID 6PW5) (Gao *et al.,* 2016; Yin *et al.,* 2017; Tang *et al.,* 2018; Paulsen *et al.,* 2015; McGoldrick *et al.,* 2019). We identified three well defined clusters (*hereafter referred to as patches*) of signature residues that are present in all representative channels, including crTRP1 from green algae (Figure 3; Supplementary Figure 2). Rat TRPV1 numbering was used throughout the manuscript to identify amino acid positions.

**Figure 3.**
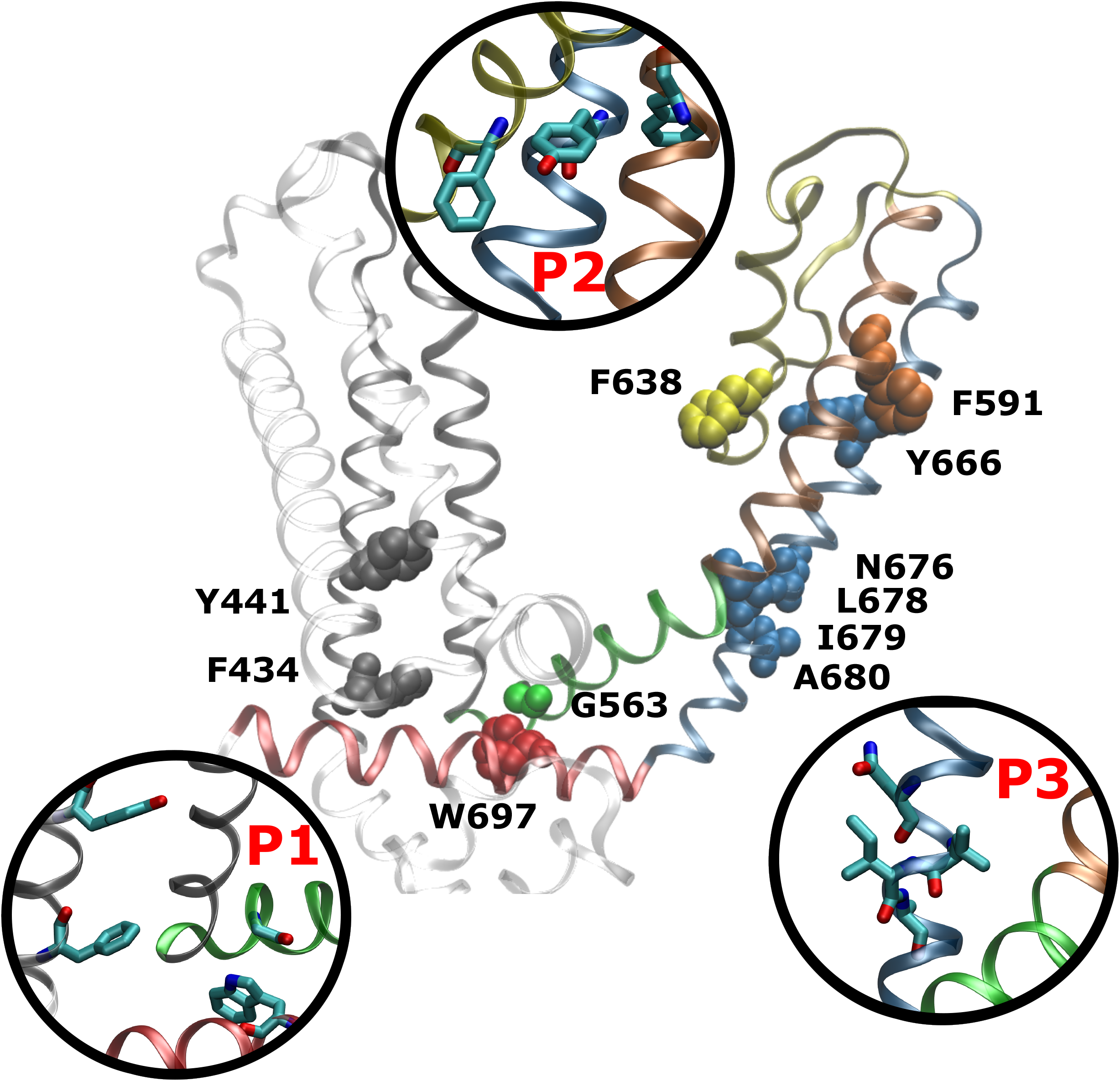
Spatial distribution of TRP channel fingerprint. Highly conserved residues are arranged in three well-defined patches, highlighted as insets and dubbed P1, P2, and P3. The structural data and residue numbering corresponds to rat TRPV1 (PDB:5IRZ). For clarity only one protomer is shown. Backbone and residues follow the code color used in Figure 2.

The first patch (P1) gathers several residues from a hotspot that has been historically linked to channel modulation. It is composed of side chains from TM1 (Phe434 [91.9%]), the TM4-TM5 linker (Gly563 [98.6%]) and the TDh (Trp697 [92.4%]) (Figure 3; *Table 2*). Initially proposed as critical for TRPV1 channel activation (Gregorio-Teruel *et al.,* 2014), these glycine at the linker and tryptophan at the TDh have been reported as a common theme in TRPs. In this context, the TDh seems to operate as an integrator, receiving information from lipids (such as PIP2), the TM4-TM5 linker that reports changes occurring at the transmembrane region and the coupling domain (CD) composed of the pre-S1 helix and a helix-loop-helix (HLH) motif. The high conservation of Phe434 is underscored by the importance of the CD array that couples with all the rest of the cytoplasmic features (Garcia-Elias *et al.,* 2015; Romero-Romero *et al.,* 2017; Hofmann *et al.,* 2017; Yang *et al.,* 2018; Hilton *et al.,* 2019; Yuan, 2019; Zubcevic *et al.,* 2019; Cao, 2020).

**Table 2.** Summary of structural-functional studies and reported effects of site directed mutagenesis in signature residues. First column indicates the equivalent signature residue in the rTRPV1 sequence. Second column indicates the channel member studied. Third column describes the amino acid identity or mutation, and fourth column corresponds to the type of study used to determine functional effects. MDS: Molecular dynamics simulations, SDM: Site-directed mutagenesis. Effects and references are described in the last two columns.

| TRPV1 Position | Channel | Residue | Evidence source | Effect | Reference |
| --- | --- | --- | --- | --- | --- |
| F434 | drTRPC4 | F366 | Structure | Part of cholesterol binding pocket. | Vinayagam *et al.*,2018 |
|  | hTRPV1 | H719A | SDM | Disrupt the voltage-gated opening | Zimova *et al.*,2018 |
| F/Y441 | TRPV1 | Y441S | SDM | Non functional | Boukalova *et al.*,2013 |
|  | TRPM8 | Y745H | SDM | Critical on Menthol Sensitivity. | Bandell *et al.*,2006 |
|  |  | Y745H | SDM | Critical on inhibition SKF96365-mediated of Cold- and voltage- activation, but just partially on other inhibitor | Malkia *et al.*,2009 |
| G563 | rTRPV1 | G563S/C | SDM | Gain of Function | Boukalova *et al.*,2010 |
|  |  | G563S/A | SDM | Gain of Function, Inhibition by proton of Max current induced by capsaicin | Boukalova *et al.*,2013 |
|  | mTRPV1 | G564S | SDM | Gain of Function | Duo *et al.*,2018 |
|  | rTRPV3 | G573S/C | SDM | Gain of Function | Xiao *et al.*,2008 |
|  |  | G573S/C | SDM | Gain of Function, Olmsted Syndrome | Lin *et al.*,2012 |
|  | mTRPC4/5 | G503S/<br>G504S | SDM | Gain of Function | Beck *et al.*,2013 |
|  | TRPC3 | G552 | Structure | Coupled W673 from TRP domain | Fan *et al.*,2018 |
|  | TRPCh | G552 | Structure | Coupled W673 from TRP domain | Fan *et al.*,2018a |
| F/Y591 | rTRPV1 | F591 | MDS | Part of the vanilloid binding pocket | Elokely *et al.*,2015 |
|  |  | F591A | SDM | Low Capsaicin response, non response to pH and not RTX binding | Ohbuchi *et al.*,2016 |
| | hTRPM4 | Y944 | Structure | Forming face to face π-stack with F1027 on TM5 | Duan *et al.*,2018 |
| F/Y638 | rTRPV1 | F638A | SDM | Gain of Function, NMDG/Na selectivity rise | Munns *et al.*,2015 |
|  |  | F638W | SDM | Enhanced the sensitivity to the acylpolyamine toxins AG489 and AG505 | Kitaguchi *et al.*,2005 |
|  | TRPM8 | Y908A/W | SDM | Not responsive to Cold and Menthol but responsive to Icilin | Bidaux *et al.*,2015 |
|  |  | Y908F | SDM | Totally responsive to Cold and Menthol and Icilin | Bidaux *et al.*,2015 |
|  | TRPC4 | F572 | Structure | Stabilizes the pore through an hydrophobic contact with neighbour protomer | Vinayagam *et al.*,2018 |
|  | mTRPC5 | F576A | SDM | Non-functional, dominant negative | Strübing *et al.*,2003 |
|  | hTRPA1 | F909A | SDM | Affect different agonists and antagonists responses | Chandrabalan *et al.*,2019 |
|  |  | F909T | SDM | Abolish the A-967079-inhibition of AITC-evoked response | Paulsen *et al.*,2015 |
| Y/F666 | TRPV1 | Y666A | SDM | Non-functional (present in membrane) | Susankova *et al.*,2007 |
|  | mTRPV3 | Y661C | SDM | Not responsive to Temp, but responsive to agonist (2-APB and Camphor) | Grandl *et al.*,2008 |
|  | hTRPV4 | Y702L | SDM | Not responsive to Temp, Agonist and Swelling | Klausen *et al.*,2014 |
|  | hTRPM6 | Y1053C | SDM | Causes hypomagnesemia with secondary hypocalcemia, Decreased Current amplitude in heterologous expression in HEK293 | Lainez *et al.*,2013 |
| | hTRPM4 | F1027 | Structure | Forming face to face π-stack with Y944 on TM5 | Duan *et al.*,2018 |
|  | hTRPA1 | F909A | SDM | Affect different agonist responses | Chandrabalan *et al.*,2019 |
| N676 | rTRPV1 | N676 | MDS | Gating relies on the rotatory motion of N676 | Kasimova *et al.*,2018 |
|  |  | N676A | SDM | Non-functional (present in membrane) | Kasimova *et al.*,2018 |
|  |  | N676F | SDM | Not responsive to Temp and Agonist (Cap/RTX) and reduced response to pH | Kuzhikandathil *et al.*,2001 |
| L678 | rTRPV1 | L678A | SDM | Low response to Agonist (Cap) and Temp, but normal response to co-treatment | Susankova *et al.*,2007 |
|  |  | L678P | SDM | Not responsive to Temp and Agonist (Cap/RTX) and reduced response to pH | Kuzhikandathil *et al.*,2001 |
|  | TRPV3 | L768F | SDM | Olmsted Syndrome and Erythromelalgia (gain of function) | Duchatelet *et al.*,2014 |
|  | TRPC3 | L654 | Structure | Constriction site in the lower region of the pore. | Fan *et al.*,2018a |
| I679 | rTRPV1 | I697 | Structure | Constriction site in the lower region of the pore. | Liao *et al.*,2013 |
|  |  | I697 | Structure | Constriction site in the lower region of the pore. | Cao *et al.*,2013 |
|  |  | I697 | Structure | Constriction site in the lower region of the pore. | Gao *et al.*,2016 |
|  |  | I697 | Structure | Constriction site in the lower region of the pore. | Chugunov *et al.*,2016 |
|  |  | I697 | Structure | Constriction site in the lower region of the pore. | Susankova *et al.*,2007 |
|  | mTRPM4 | I1036 | Structure | Constriction site in the lower region of the pore. | Guo *et al.*,2017 |
|  | hTRPM4 | I1040 | Structure | Constriction site in the lower region of the pore. | Autzen *et al.*,2017 |
|  | drTRPC4 | I617 | Structure | Constriction site in the lower region of the pore. | Vinayagam *et al.*,2018 |
|  | TRPV4 | I715N | SDM | Hydrophobic single-residue gate. Higher resting currents. | Zheng *et al.*,2018 |
|  | TRPC4 | I617N | SDM | Hydrophobic single-residue gate. Higher resting currents. | Zheng *et al.*,2018 |
|  | TRPM8 | V976S | SDM | Hydrophobic single-residue gate. Higher resting currents. | Zheng *et al.*,2018 |
| A680 | TRPV1 | A680 | MDS | Change of Solvation | Chugunov *et al.*,2016 |
| | TRPV4 | A716S | SDM | Not responsive to agonists (4αPDD, Hypotonicity and AA), cause SMD Kozlowski type, and Metatropic Dysplasia | Krakow *et al.*,2009 |
|  | hTRPVA1 | G955A | SDM | Slower inactivation rate. Lower rectification rates. | Benedikt *et al.*,2009 |
|  |  | G958R | SDM | Inward-rectifier, constitutively active at resting potential, and impaired response to AITC | Benedikt *et al.*,2009 |
| W697 | rTRPV1 | W697 | Structure | It forms a hydrogen bond with the main chain carbonyl oxygen of F559 at the beginning of the S4-S5 linker | Liao *et al.*,2013 |
|  |  | W697A | SDM | Low Response to Cap/Em | Valente *et al.*,2008 |
|  |  | W697x | SDM | Low Response to Cap/Em, Affect allosteric activation | Gregorio-Teruel *et al.*,2014 |
|  | TRPV3 | W692G | SDM | Gain of Function, Olmsted Syndrome | Lin *et al.*,2012 |
|  | TRPV4 | W733R | SDM | Gain of Function, limited agonist response, and not inactivation to long depolarization | Teng *et al.*,2015 |
|  | TRPC3 | W673 | Structure | It is extensively coupled with the S4-S5 linker through interactions with G552 | Fan *et al.*,2018a |
|  | TRPC4 | W674 | Structure | Coupled with the S4-S5 linker through interactions with G553 and P546 on TM4 | Fan *et al.*,2018 |

**Table 3.** Summary of mutation effects of residues from the AC or the conserved TM4 residue connecting TM4 with TM5 in the literature. First column indicates the equivalent signature residue in the rTRPV1 sequence. Second and third columns indicate the channel studied and the specific mutation introduced, respectively. Fourth column corresponds to the effect of the mutation and/or proposed function.

| TRPV1<br>Position | Channel | Mutation | Effect | Reference |
| --- | --- | --- | --- | --- |
| F/Y444 | TRPV1 | Y444S | Non-functional | Boukalova *et al.*,2013 |
|  | mTRPM3 | Y885T | Impaired non-canonical current induced by pregnenolone sulphate + clotrimazol | Held *et al.*,2018 |
| F448 | TRPV1 | F448L | Decreased pH response but maintain all Cap responsiveness | Boukalova *et al.*,2013 |
|  | mTRPM3 | Y888T | Similar to *wt* response to pregnenolone sulphate + clotrimazol | Held *et al.*,2018 |
| Y/F554 | TRPV1 | Y554A | Non-functional (Cap 10μM, -70 to 200mV, 48°C) | Boukalova *et al.*,2010 |
|  |  | Y554F | Normal responsiveness | Boukalova *et al.*,2010 |
|  |  | Y554A | Not responsive to pH, Cap and RTX | Elokely *et al.*,2015 |
|  |  | Y554A | Increased sensitivity and affinity to 2-APB | Singh *et al.*,2018 |
|  | TRPV2 | Y514A | Increased sensitivity and affinity to 2-APB | Singh *et al.*,2018 |
|  | TRPV3 | Y564A | Increased affinity to 2-APB | Singh *et al.*,2018 |
| Y/F555 | TRPV1 | Y555S | Non-functional (Cap 10μM, -70 to 200mV, 48°C) | Boukalova *et al.*,2010 |
|  |  | Y555F | Normal responsiveness | Boukalova *et al.*,2010 |
| W549 | rTRPV1 | W549A | Not responsive to Cap (and RTX and others) and pH | Ohbuchi *et al.*,2016 |
|  | rTRPV2 | W549A | Interaction with vanillyl moiety of RTX or Capsaicin | Gavva *et al.*,2004 |
|  | hTRPV4 | W568A | Impaired responsiveness to heat and agonists (4α-PDD and BAA); but responsive to swelling and endogen lipids | Vriens *et al.*,2007 |
|  | mTRPM3 | W982R | Abolished non-canonical current induced by pregnenolone sulphate + clotrimazol | Held *et al.*,2018 |
|  | mTRPM3 | W982F | Similar to *wt* response to pregnenolone sulphate + clotrimazol | Held *et al.*,2018 |
|  | hTRPA1 | Y840F | Reduced potency of ligand (GNE551) | Liu *et al.*,2021 |
|  | hTRPA1 | Y840W/<br>H/L/A | Completely abolished potency of ligand (GNE551) | Liu *et al.*,2021 |
|  | hTRPA1 | Y840A | Impaired response to AITC and almost abolished to β-Eudesmol | Ohara *et al.*,2015 |

The second patch (P2) is located at the selectivity filter region and is composed of three phenyl group residues, *i.e.*, Phe591 [Φ 97.4%], Phe638 [Φ 97.2%], and Tyr666 [Φ 98.7%], located at both the TM6 helix and the pore helix (Figure 3). A glycine residue that forms part of the selectivity filter/upper constriction in TRPs is also highly conserved (87.9%) but slightly off to the 90% threshold (Figure 2). The high conservation of these residues within the selectivity filter contrasts with the high variability observed by us and others at the pore region (Huffer *et al.,* 2020).

A third patch (P3) is localized at the lower portion of the pore and is composed of a well-studied set of residues forming the lower gate (Asn676 [95.0%], Ile679 [90.4%], Ala680 [94.4%]) as well as Leu678 [95.4%] that is facing the interface between TM5 and TM6 (Palovcak *et al.,* 2015) (Figure 3).

Finally, we identified a prevalent aromatic side chain (Tyr441 [76.6% Tyr + 15.2% Phe]), located at the middle of the TM1 helix. This residue was not observed in close contact to any other signature residue (Figure 3).

From the structural data, the most conserved interaction between signature residues is between the glycine (Gly563) in the TM4-TM5 linker and the tryptophan (Trp697) at the TDh. This observation is further supported by our evolutionary coupling analysis showing that the highest score for putative interactions-within the region we studied- are precisely those established between the TM4-TM5 linker and the TDh (Supplementary Figure 3). The analysis also showed interactions linking the lower portion of TM2 to the end of the TDh. Such interaction cannot be explained by direct contact between TM2 and TDh. Thus, it would not be unreasonable to suggest that this interaction between TM2 and the TDh involves an additional linker molecule such as PIP_2_ or other lipid binding to this region (Poblete *et al.,* 2014; Yin *et al.,* 2017; Yazici *et al.,* 2021; Hughes *et al.,* 2018)). Interactions between the channel and the membrane at the cytosol-membrane interface is emerging as a common theme in TRPs. Under this view, different parts of the CD/TDh coupling mechanism are tuned by the differences in binding of membrane lipids and/or canonical ligands (reviewed in Zubcevic, 2020). Moreover, a calcium binding site located at the intracellular linker has been observed in several TRP channel structures and associated with channel function (Brauchi & Rothberg, 2020). To our knowledge, calcium-dependent modulation of the putative TM2-lipid-TDh interaction has not been explored in detail.

Series of independent studies had reported that mutations of residues that form part of the fingerprint -or the connecting side chains-modify or impairs channel activity, underscoring their importance in maintaining proper channel activity (Table 2). Then, we analyzed the networks of connectivity between the signature residues in channels from different subtypes. The apo and ligand-bound state conformations of prototypical channels were used to compare state-dependent changes in connectivity (*i.e.,* TRPV1/2/3/5/6, TRPC3/4/6, TRPM2/4/8 and TRPA1). As reported abundantly in the literature, the signature residues in the TM4-TM5 linker and TDh from P1 and the residues in P3 forming the lower gate experience changes in their network of connectivity (Supplementary Figure 4). On the other hand, residues in P2 appear displaced altogether as result of pore reconfiguration with just minor changes in distances between them (Supplementary Figure 4).

### A conserved aromatic core at the transmembrane region of TRPs

Four phenyl groups (*i.e.*, Phe/Tyr) were identified in specific positions within the transmembrane domain in more than 75% of the sequences. Further inspection of structural data showed that these residues are part of a larger cluster of aromatic amino acids, common to all TRP channel structures (Figure 4). A group of six to seven aromatic residues belonging to the VSLD/LBD domain (including at least one signature residue from P1), appear to form an aromatic core (AC) (Figure 4 a and b). In contrast, voltage-gated potassium and sodium channels occupy these positions with charged amino acids forming salt bridges, short aliphatic side chains, or aromatics located at the membrane-water interface (Supplementary Figure 5). Although the overall three-dimensional shape of the AC varies, it is present in all TRP subtypes, connecting three to four helices from within the VSLD/LBD, suggesting they serve as scaffold that is holding together the whole VSLD/LBD domain (Figure 4b).

**Figure 4.**
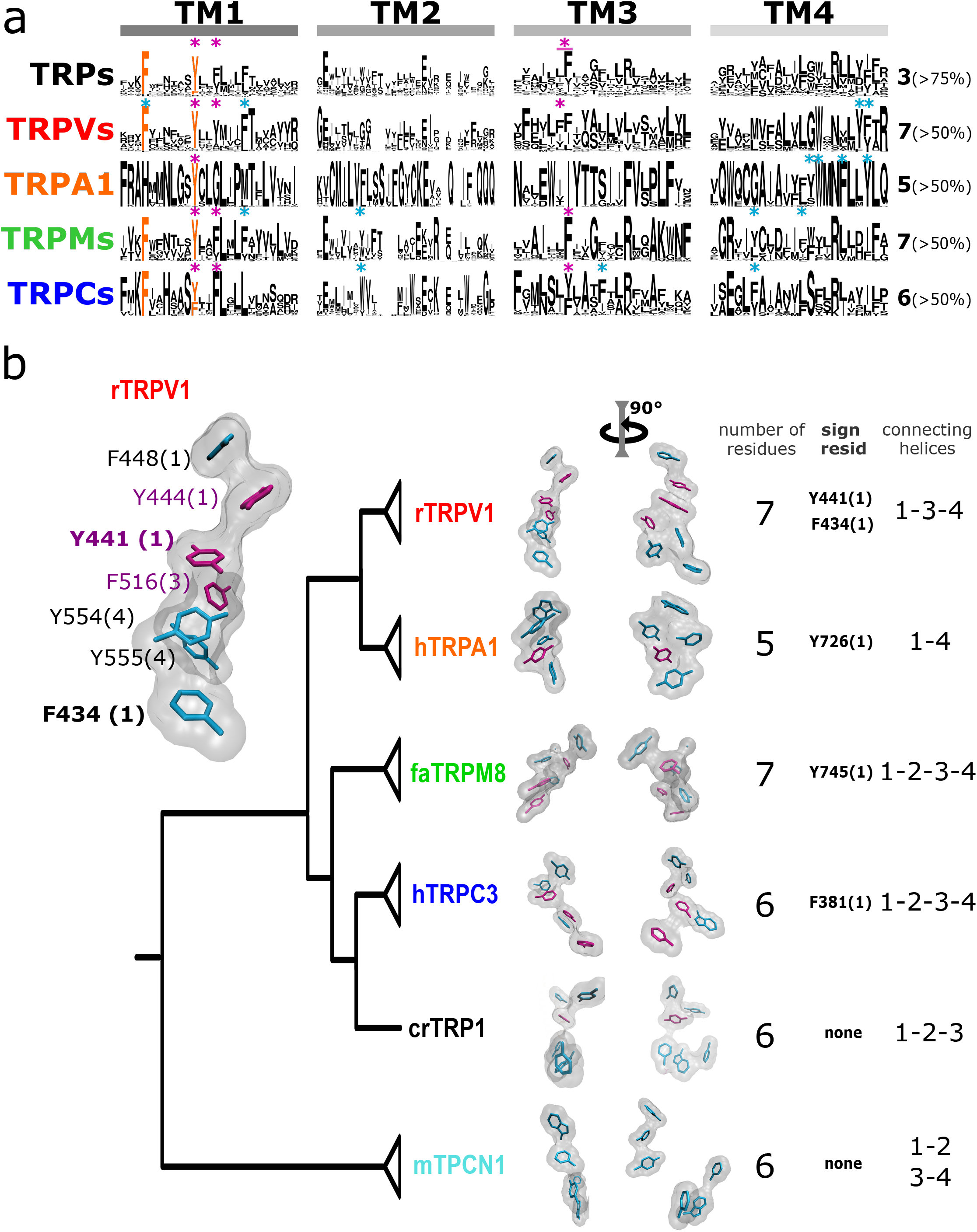
Aromatic residue distribution in LBD. All the highlighted positions comply two conditions, i.e., to be facing the core of the LBD, and to be at <4.5 Å from another aromatic residue. a) Conservation of aromatic residues on transmembrane helices 1 to 4. Orange letters: the two fingerprint aromatics in TM1. Violet asterisk: >50% of aromatic conservation within subfamilies. Cyan asterisks: >50% of aromatic conservation within the subfamily. b) Comparison between AC volumes presented next to a schematic view of the topology obtained in our phylogenetic analysis. Cyan licorice: conserved residues (>50%) in specified subfamily, Magenta licorice: conserved residue (>50%) in all families. Inset. Aromatic core in rTRPV1. The specific positions of the aromatics are indicated. Structures used: rTRPV1, PDB: 5IRZ; faTRPM8, PDB: 6BPQ; hTRPC1, PDB: 5ZBG; hTRPA1, PDB: 3J9P; crTRP1, PDB:6PW5; mTPCN1, PDB: 6C96

The AC emerges as a common theme present from crTRP1 to TRPV1-4. The most dramatic case is observed in the TRPV1-4 clade, exhibiting both the highest number of aromatics (7) and the most ordered stack (Figure 4 b). In contrast, TRPA1 shows only 5 aromatics forming the AC and a wider disposition, with only two helices connected through the aromatic network. Although the residues in TRPA1 are still in close contact, they do not form the compact stacking observed in other TRPs, including crTRP1.

The presence of these conserved aromatics next to ligand binding sites -or even forming part of it-suggests a mechanism in which the AC acts by imparting rigidity to the region and functionally linking the transmembrane helices 1-4, facilitating the translation of mechanical force from the ligand binding sites to the coupling domains that connect multiple parts of the channel including the TDh, cytoplasmic CD, and preTM1. Minimal or no displacement among these aromatics was observed by comparing the apo and ligand-bound structures from different TRP members (Supplementary Figure 4). This suggests to us that their role might be related to imposing rigidity to the VSLD. Rigidity that might be affected by ligand binding as part of a process that has been tuned during evolution in the different subfamilies. Moreover, consistent with the notion that TPCN channels are close relatives of the TRP family, the presence of a group of 6 aromatics facing inside the VSLD is conserved in TPCN’s domain 1 and absent in domain 2 that is devoid of aromatics, resembling Nav channels. Unlike TRPs, this AC in TPCN channels looks less cohesive, or “disconnected” (Figure 4b).

### Conserved residues at the interaction between subunits

TRP channels show a domain-swapped configuration. That is, the VSLD/LBD of one subunit appears in close contact with its pore domain (PD) through the protein backbone and, at the same time, in close contact with the PD of the neighboring subunit. The coupling between the VSLD and the pore domain from different subunits of the tetramer is one topic that has been poorly studied in TRP channels and not well understood in VGICs (Carvalho-de-Souza & Bezanilla, 2019; Shem-Ad *et al.,* 2013; Neale *et al.,* 2003).

By inspecting the structural data, we found a conserved interaction formed by a residue at the middle of TM4 and a residue that is consistently preceding a signature residue in TM5 (Phe591 in rTRPV1; Figure 5a and b). Although the nature of the interaction varies, it is present in all surveyed structures. Such interaction would put in direct contact TM helices 4 and 5 from different subunits (*i.e.* the VSLD and the pore). By extension, this interaction would communicate patches P1 and P2 from different subunits through the aromatics running alongside the VSLD/LBD. The high conservation of these residues and their relative positions within the structure suggests a common mechanism in TRPs where the selectivity filter (P2) would be functionally connected to the ligand binding site located on a neighboring transmembrane region.

**Figure 5.**
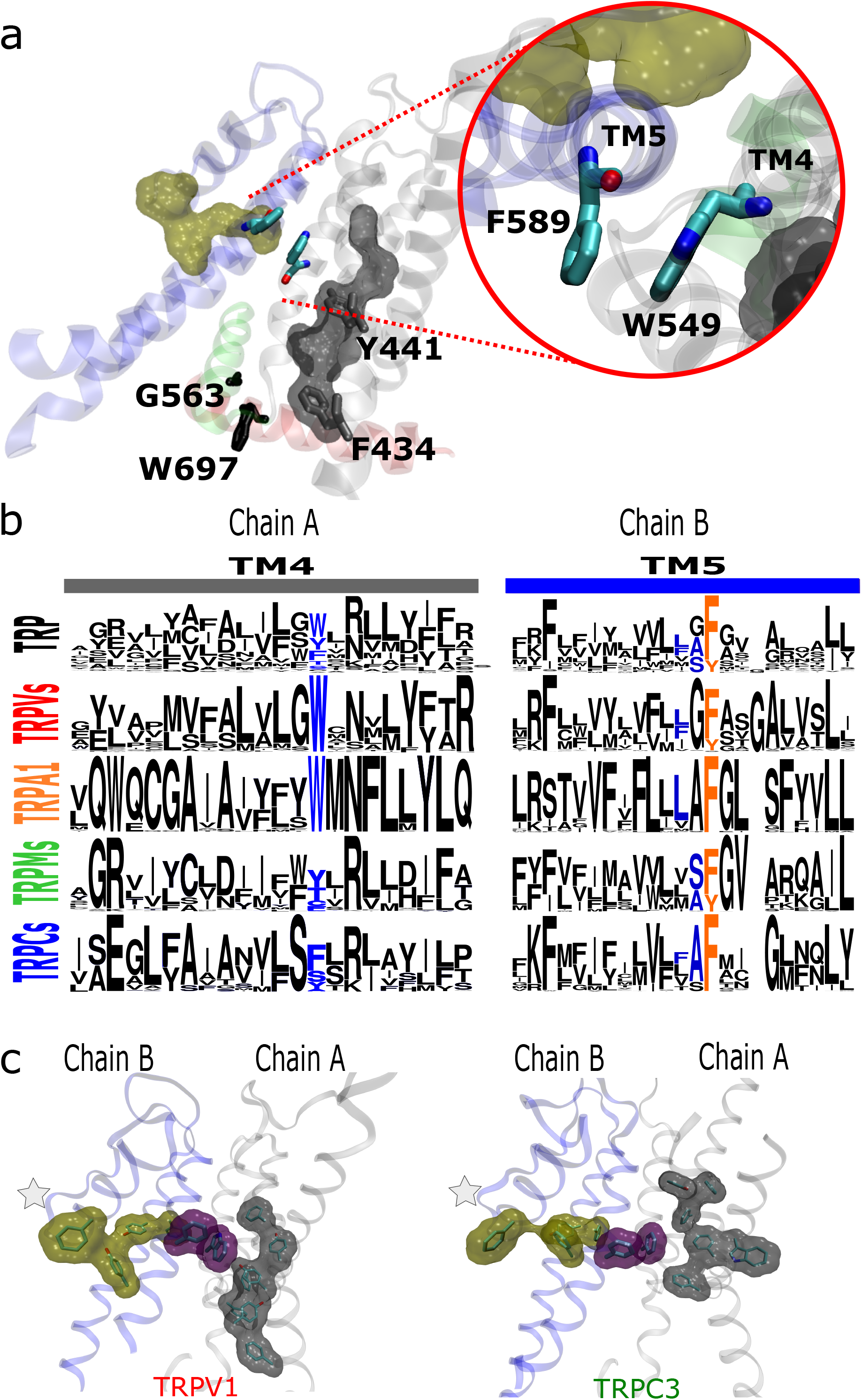
The AC connects to aromatics in P2 from neighboring subunits. a) A conserved intermolecular connection between residues (licorice) in helices at opposite faces to the AC (gray surface) and P2 (yellow surface). Signature residues from P1 are depicted in gray (F434 and Y441) and black (G563, and W697). Inset: Upper view of residues establishing the inter-subunit interaction (PDB:5IRZ). b) Residues connecting TM4 and TM5 from neighboring subunits align on the same position in TM4 and at one or two positions from the 4th fingerprint residue in TM5. c) Lateral view of the interaction between LBD and PD from different subunits in the most distant members of the TRP clade.

## DISCUSSION

In the present work we studied the conservation and frequency distribution of amino acids within the transmembrane of TRP channels and flanking regions, trying to conciliate structural and evolutionary contexts (Figure 6). A highly conserved set of residues are located and grouped at strategic positions within the channel’s transmembrane region, defining a fingerprint for GI-TRPs. Moreover, a conserved network of interactions connects them directly or indirectly.

**Figure 6.**
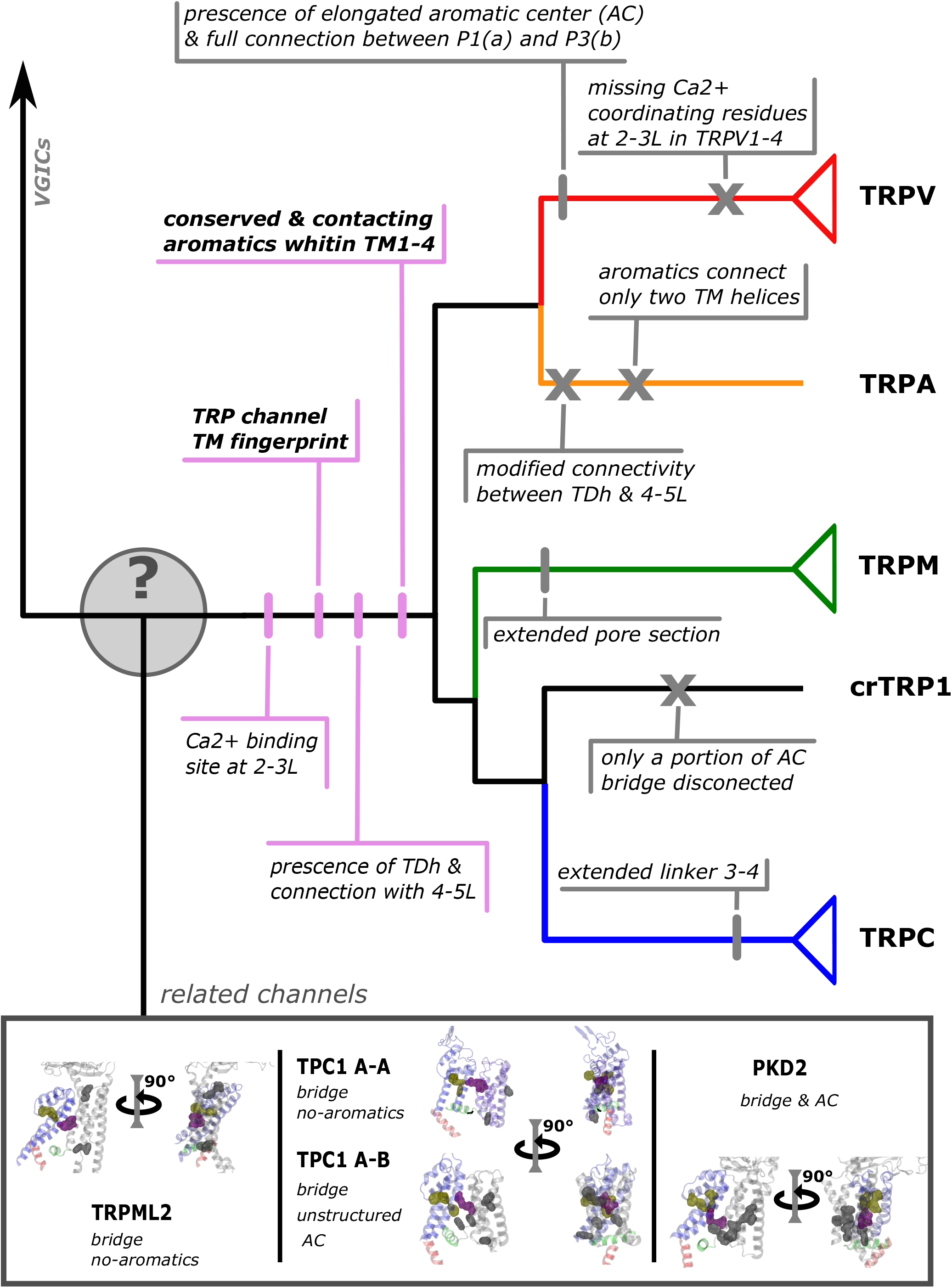
Conserved modif ications observed in TRP proteins. The different channels studied in this work are presented next to a schematic view of the topology obtained in our phylogenetic analysis. Unique TRP features are highlighted in pink shades. Other features that appeared segregated by families are highlighted in grey shades.

From the three patches described in this article, the aromatics forming patch 2 were not described in the literature as such. Nevertheless, previous studies showed the importance of the residues composing this patch. Phe591 is described as forming part of the higher, wider region of the capsaicin pocket binding in TRPV1 channels (Elokely *et al.,* 2015) and mutations at Tyr638 and Tyr666 have obvious effects on channel activity. Specifically, Tyr666Ala is not functional in TRPV1 (Susankova *et al.,* 2007), the equivalent mutant Tyr661Cys in TRPV3 is not activated by temperature, but still responds to agonist (Grandl *et al.,* 2008), and the Tyr702Leu equivalent mutant in TRPV4 has significantly reduced responses to agonists, temperature, and mechanical stimulation (Klausen *et al.,* 2014). Moreover, mutations at equivalent positions to rTRPV1 Tyr638 in channels from different subfamilies elicit a wide range of effects from gain-of-function to dominant-negative phenotypes (Munns *et al.,* 2015; Kitaguchi & Swartz, 2005; Bidaux *et al.,* 2015; Strübing *et al.,* 2003; Chandrabalan *et al.,* 2019; Vinayagam *et al.,* 2018; Paulsen *et al.,* 2015). Noteworthy, Bidaux *et al.,* (Bidaux *et al.,* 2015) demonstrated that mutations at this position (Tyr908 in TRPM8) to Ala or Trp abolish the response to temperature and menthol but not to icilin, yet mutation to Phe keeps the channel fully functional. As shown here, the three aromatics forming P2 always need a non-conserved fourth hydrophobic residue to connect them all. By comparing apo and ligand bound structures, P2 consistently seems to translate as if it were a near-rigid-body, moving as a whole compact structure. Our analysis suggests that the function of this patch might be associated with the communication between the VSLD/LBD and the selectivity filter from different subunits.

Residues on the other two patches have been recurrently studied in the literature. In particular, residues of patch 3 have been characterized as shaping the lower gate of TRP channels and to participate in stabilizing the transition between α- and π-helix types during opening (Palovcak *et al.,* 2015; Kasimova *et al.,* 2017; Kasimova *et al.,* 2018).

### The interaction between signature residues and the aromatic core

Two of the residues in the P1 patch have been proposed as fundamental to the interaction between the TDh and the TM4-TM5 linker, acting as an allosteric integrator (Taberner *et al.,* 2014; Sierra-Valdez *et al.,* 2018; Romero-Romero *et al.,* 2017; Gregorio-Teruel *et al.,* 2014; Zhao *et al.,* 2020). A third residue in P1 corresponds to a conserved Phe at the lower end of TM1 that is always connected to the other elements of the patch. Separated by 2 intermediate interactions, TRPM8 displays the largest distance between these two components of patch P1. Notably, the gap observed between these two components has been associated to the binding site of menthol (Bandell *et al.,* 2006; Malkia *et al.,* 2009). Moreover, the menthol analog WS-12 binds to Tyr745 (TM1), and sits close to Tyr1004 (TDh), seemingly reinforcing the connection between the VSLD/LBD and the end of the TDh in TRPM8 (Yin *et al.,* 2019).

There is only one signature residue that is consistently forming part of the aromatic core (*i.e.*, Tyr441 in TRPV1). Mutations to Ser of Tyr441 in TRPV1 generates non-functional channels (Boukalova *et al.,* 2013). The equivalent residue in TRPM3 channels presents impairments in response to agonists when mutated from Tyr to Thr, and a Tyr to His mutation in TRPM8 channels has been shown critical to both menthol activation and the inhibitory effect of the small molecule SKF96365 (Bandell *et al.,* 2006; Malkia *et al.,* 2009). This coincided with similar phenotypes observed by mutating other residues belonging to the AC. This is the case of Tyr444 that in TRPV1 generates non-functional channels (Boukalova *et al.,* 2013) and the double mutation Y885T / W982R in TRPM8 that presented altered responses to agonists and temperature (Held *et al.,* 2018).

When comparing the disposition of the aromatic core in detail, it appears obvious they have different distributions in the different TRP channel families. However, certain generalizations can be drawn. The AC appears associated with P2 at the selectivity filter and disconnected from P1 in most families. Notably, in TRPVs the AC extends down to reach the lower aromatic residues of P1 making a direct connection between the AC and the TDh / TM4-TM5 linker. Hence, a P2-AC-P1 continuum can be observed in the TRPV subgroup. Interestingly, from the structures obtained in the presence of agonists it becomes evident that these agonists bridge the AC and P1 (e.g., WS-12 in TRPM8 and GFB-9289 in TRPC4; PDBIDs 6NR2 and 7B16, respectively; Yin *et al.,* 2019; Vinayagam *et al.,* 2020). The disposition of the components of P1 (*i.e.*, TM1;TM4-TM5linker;TDh) and differences in the AC suggest similar but nonidentical coupling strategies among GI-TRPs.

The presence of an aromatic core, connecting the transmembrane helices of the ligand binding pocket and communicating critical modulatory regions that are far apart in the structure (*e.g.,* the selectivity filter and the TDh) appears as a modular connector that has been subject to variations throughout evolution. This suggests a common aspect of TRP channel mechanics that remains largely unexplored. Our line of reasoning also suggests that the observed rearrangements of the selectivity filter during activation are likely linked to inter-subunit interactions, likely modulated by the AC. At the same time, it implies that certain ligands might have the ability to modulate the conformation of the selectivity filter - and by extension the extracellular linkers-without the need of an open gate conformation.

## METHODS

### Amino Acid sequences, alignment, and phylogenetic analyses

Protein sequences corresponding to TRPA1, TRPC1, TRPC3-7, TRPM1-8, TRPML1-3, TRPP1-3, and TRPV1-6 from human (*Homo sapiens*), TRPC2 from the house mouse (*Mus musculus*), NompC from the fruit fly (*Drosophila melanogaster*) and TRPY1 from the brewing yeast (*Saccharomyces cerevisiae*) were retrieved from the International Union of Basic and Clinical Pharmacology (IUPHAR) database. The transmembrane regions of these sequences were used as queries in blastp searches (Altschul *et al.,* 1990) against the proteomes of *Chlamydomonas reinhardtii*, *Volvox carteri, Coccomyxa subellipsoidea*, *Micromonas pusilla*, *Dictyostelium discoideum*, *Dictyostelium purpureum, Leishmania infantum*, *Leishmaniamajor, Leishmania mexicana, Paramecium tetraurelia*, and *Trypnosoma cruzi* to search for putative TRP channels. Putative channels were selected based on the frequency of hits to the query sequences relative to human, friut fly, and yeast. This was followed by reciprocal blastp searches (E-value < 1e), and a final inspection for the presence of the TRP domain. To investigate the phylogenetic position of these candidate TRP channels we retrieved protein sequences corresponding to TRPVs, TRPA1, TRPMs, TRPN1, and TRPCs in representative species of all main lineages of amniotes (Supplementary Table S1) from the Orthologous MAtrix project (OMA) (Altenhoff *et al.,* 2021). Thus our final dataset included 844 protein sequences from *bona fide* TRP channels from amniotes in addition to 24 sequences from unicellular organisms. Amino acid sequences were aligned using MAFFT v.7 (Katoh *et al.,* 2017) allowing the program to choose the alignment strategy (FFTNS2). We used the proposed model tool of IQ-Tree v.1.6.12 (Minh *et al.,* 2020) to select the best-fitting model of amino acid substitution (JTT+G4). We used the maximum likelihood method to obtain the best tree using the program IQ-Tree v1.6.12 (Minh *et al.,* 2020). We carried out 3 independent runs to explore the tree space, and the tree with the highest likelihood score was chosen. We assessed support for the nodes using the ultrafast bootstrap routine as implemented in IQ-Tree v1.6.12 (PMID: 29077904). Human (*Homo sapiens*) and chicken (*Gallus gallus*) Potassium voltage-gated channel subfamily A member 2 (KCNA2), Sodium Voltage-Gated Channel Alpha Subunit 8 (SCN8A) amino acid sequences were included as an outgroup.

### MSA database

474 extra sequences from the Uniprot protein database were added to the pool of sequences rescued from the OMA server and then realigned in MAFFT. The region corresponding to the transmembrane was identified, and the rest of the sequence was removed, leaving the section from the last portion of the pre-TM1 region to the last residue of the TRP helix (using rTRPV1 as reference). A second round of sequence alignment in MAFTT (FFTNS1 strategy) was performed and manually refined to minimize gaps. The resulting database contains 1311 sequences, 837 selected from OMA database and identified unicellulars and 474 from Uniprot, including sequences annotated as TRP or TRP-like channels and several uncharacterized protein sequences.

### Coevolution analysis

Coevolution scores were calculated using asymmetric pseudo-likelihood maximization direct coupling analysis algorithm (aplmDCA), (Ekeberg *et al.,* 2013). This algorithm finds the approximate parameters of the maximum entropy probabilistic model consistent with selected MSA statistics (univariate and bivariate frequency distributions). Default parameters were used for field and coupling regularization and sequence reweighting (lamba_h = 0.01, lambda_j = 0.01, theta = 0.1).

### Structures

The structures used were rTRPV1, PDB entry 5IRZ; faTRPM8, 6BPQ; hTRPC3, 5ZBG; hTRPA1, 3J9P; crTRP1, 6PW5; mTPCN1, 6C96; rKv1.2, 3LUT and hNav1.4, 6AGF. To compare the apo vs ligand-bound/activated channel structures, the PDB files used were PDB:5IRZ -apo-, PDB:5IRX -RTX +DkTx-), *Ficedula albicollis* TRPM8 (PDB:6BPQ –apo-; PDB: 6NR2 -Ws-12 + PIP2-) and *Danio rerio* TRPC3 (PDB: 6G1K –apo-; PDB:7B16 -GFB-9289-).

### Figure preparation

To visualize the identities and gap patterns on the MSA, images were exported using Jalview 2.11.1.3 (Waterhouse *et al.,* 2009). The sequence logos show the distribution of amino acid residues at each position in the regions of interest, and were generated using WebLogo version 3 (Crooks, 2004). The structural figures were generated using VMD 1.9.2 (Humphrey *et al.,* 1996). Direct interactions between residues were identified using a distance threshold of 4 Å, except for the pi-pi interactions, where a 5 Å threshold was used (Piovesan *et al.,* 2016).

## Acknowledgements

We acknowledge Dr. Gonzalo Riadi from Universidad de Talca for providing computer support. JCO acknowledges the Integrative Biology Group members, Universidad Austral de Chile, for their constant support, scientific enthusiasm, and creative feedback. This work was supported by Fondo Nacional de Desarrollo Científico y Tecnológico from Chile (FONDECYT 1191868) to SB, (FONDECYT 1210471) to JCO, and ANID-Millennium Science Initiative Program #NC160011 (SB and JCO).

## Author contribution

SEB, CKC, and DCB designed experiments and analyzed data. VC, KZ and JCO designed the taxonomic sampling, retrieved amino acid sequences and performed phylogenetic analyses. DCB and SEB prepared the figures. DCB, CKC, SEB and JCO wrote the manuscript.

---

**Figure S1.**
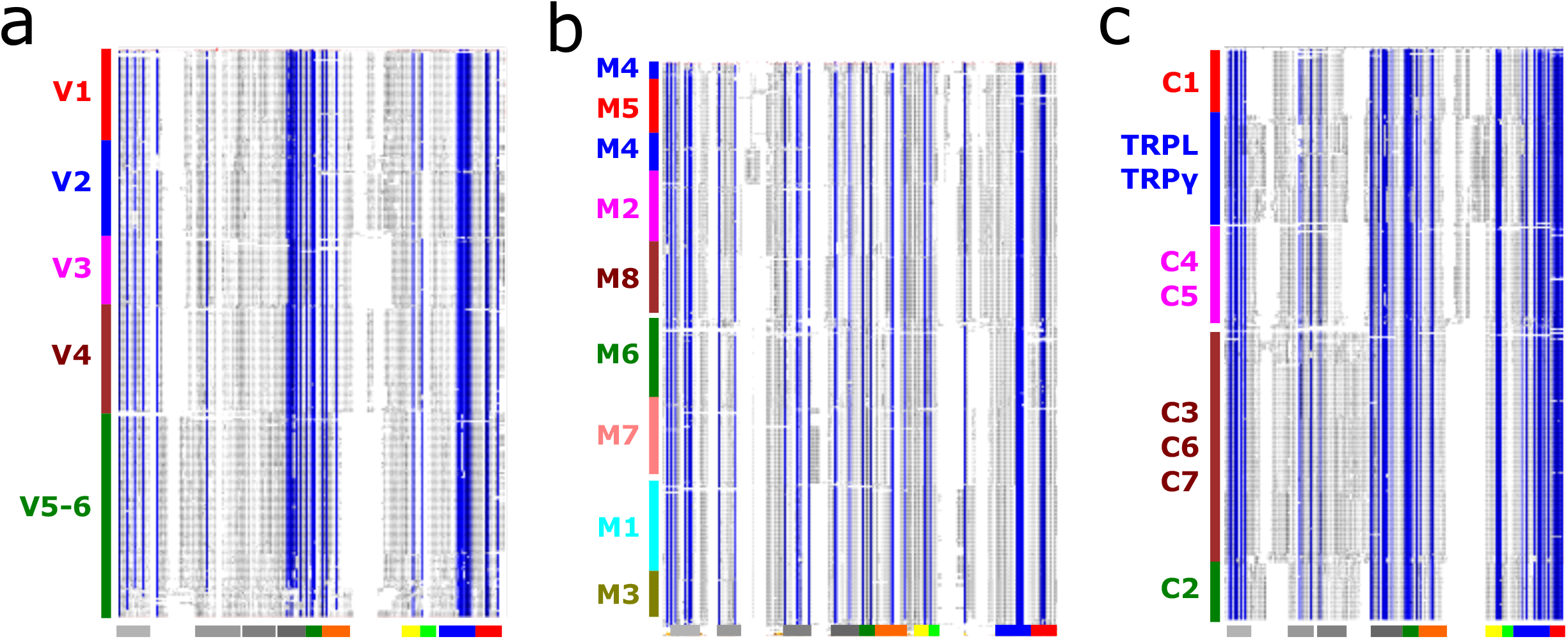
MSAs for TRP channel subfamilies V (a), M (b), and C (c). Sequences are stacked in the Y axis and progression from N- to C-terminal on the X axis. Blue vertical lines indicate residues with >90% of identity. Below each panel,structural features are shown in bars following the color code in Fig. 2.

**Figure S2.**
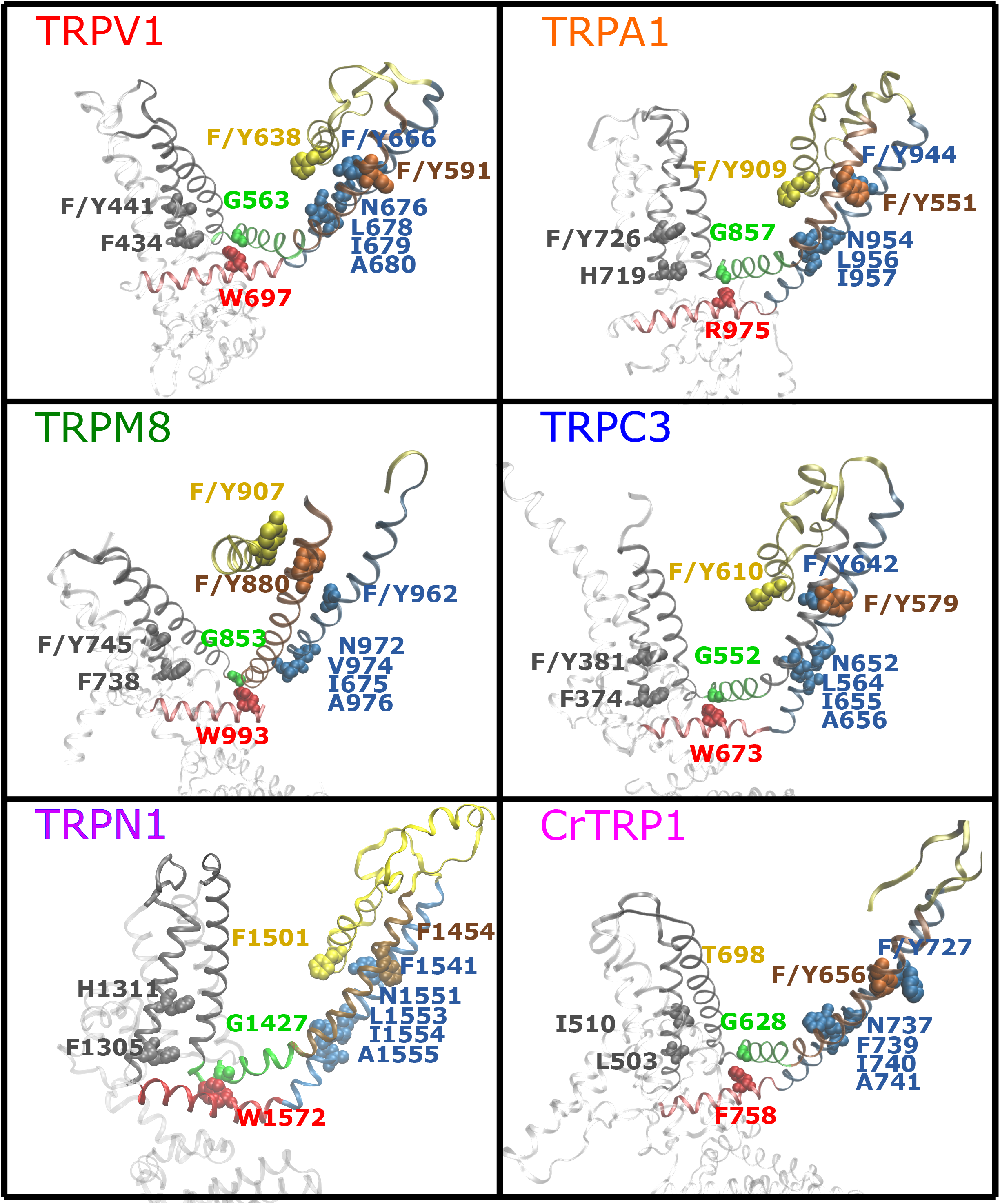
TRP fingerprint. Conserved residues and the forming patches are presented for a member of each TRP subfamily. Residues are colored according to the color cod e in Fig. 2. Structures used: rTRPV1, PDB: 5IRZ; faTRPM8, PDB:6BPQ; hTRPC3, PDB:5ZBG; hTRPA1, PDB:3J9P; CrTRP1, PDB:6PW5.

**Figure S3.**
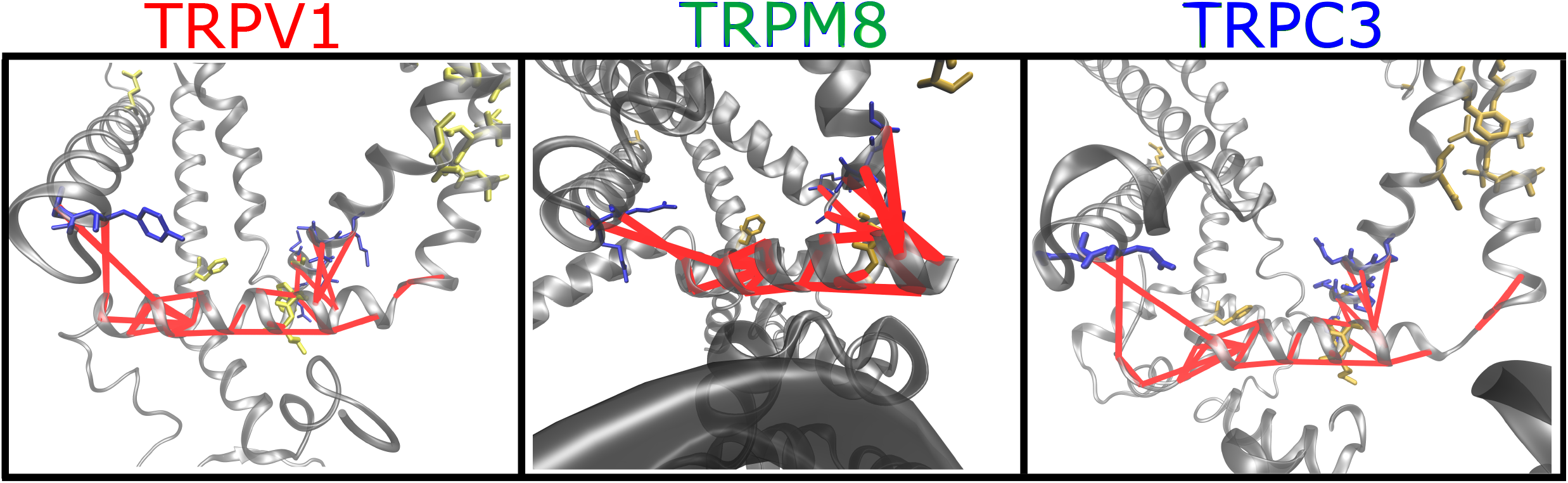
Coevolution analysis. Residue pairs with high coevolution scores (top 5%) are connected by red lines. Coevolution scores were calculated using asymmetric pseudo-likelihood maximization direct coupling analysis algorithm (aplmDCA). Signature residues are drawn in blue licorice representation and non-signature residues in yellow (rTRPV1, PDB: 5IRZ; faTRPM8, PDB:6BPQ; hTRPC3, PDB:5ZBG).

**Figure S4.**
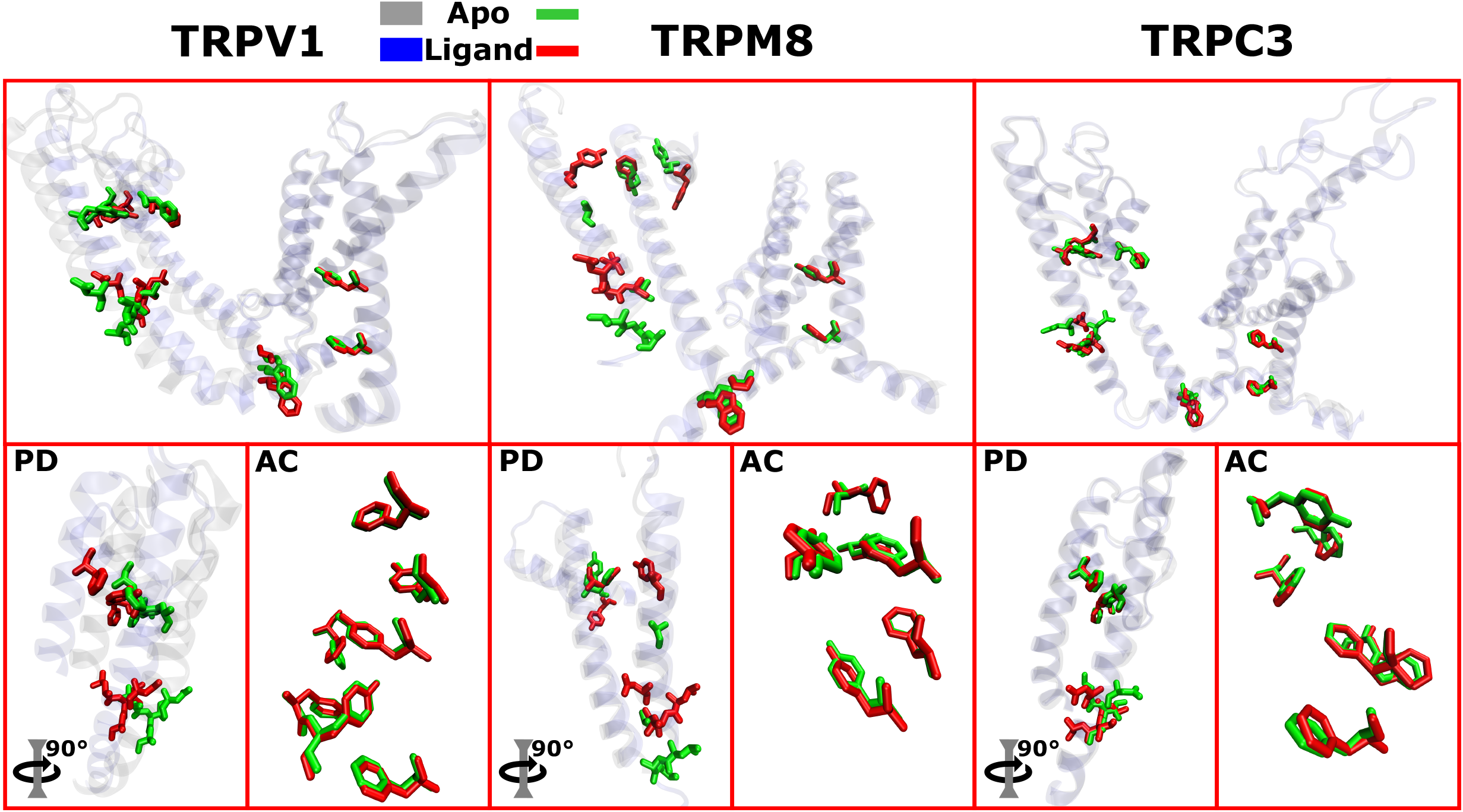
Conformation-dependent changes on spatial distribution of conserved residues. Upper boxes: Relative position of fingerprint residues in the apo(green licorices on gray ribbons) and ligand bound (red licorices on blue ribbons) structures. Bottom boxes: Relative position of residues from the pore domain (PD) and the aromatic core (AC) comparing between apo and ligand bound structures. (rTRPV1 apo, PDB: 5IRZ; TRPV1ligand-bound, PDB: 5IRX; faTRPM8 apo, PDB: 6BPQ; faTRPM8 ligand-bound: 6NR2; hTRPC3 apo, PDB: 5ZBG; hTRPC3 ligand-bound, PDB: 6CUD).

**Figure S5.**
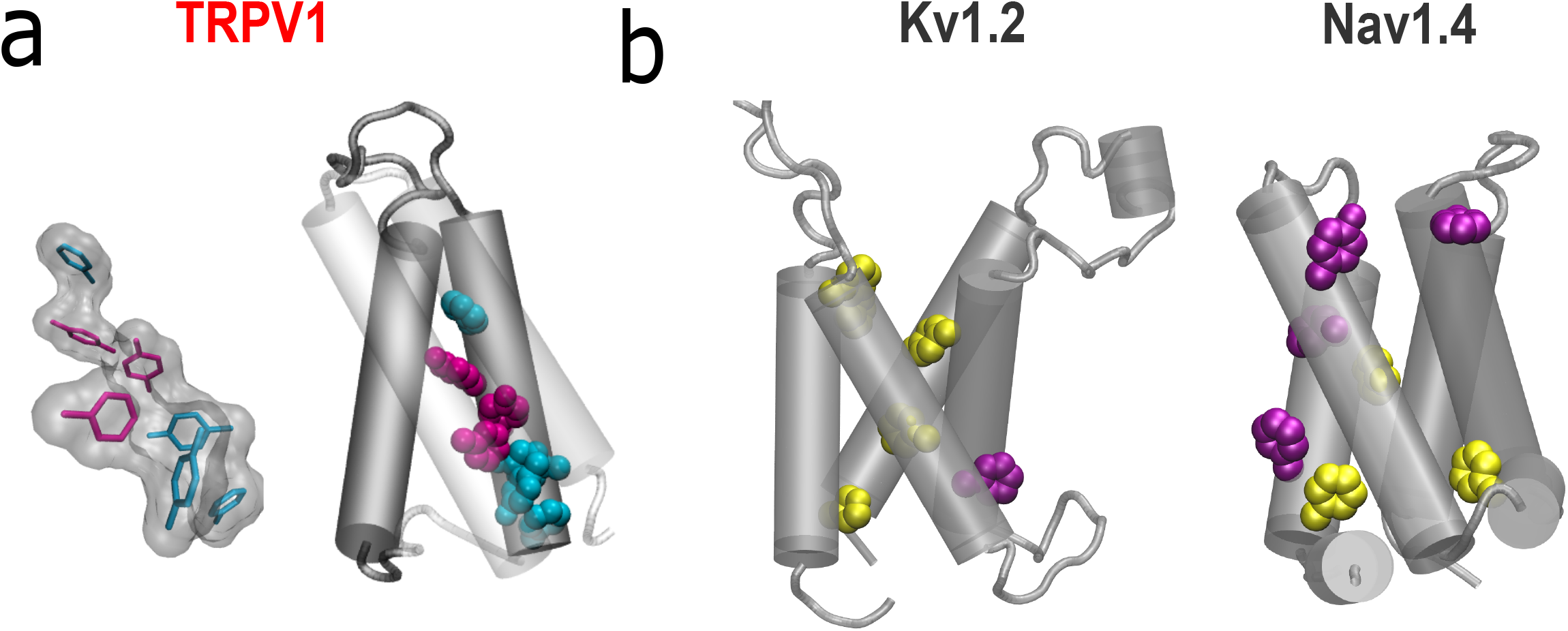
Aromatic residues in non-TRP channels. a) TRPV1 aromatic core. Cyan licorice/spheres: conserved residues (>50%) in the specified subfamily, Magenta licorice/spheres: conserved residue (>50%) within subfamilies. b) aromatic residues found in the VSD of Kv1.2 and Nav1.4. Magenta: aromatic residues aligning with the conserved aromatic in at least one TRP subfamily. Yellow: non-aligned aromatic residues present in VSD. Structures: rTRPV1 apo, PDB: 5IRZ; rKv1.2, PDB:3LUT and hNav1.4, PDB:6AGF.

